# Elucidating the patterns of seed-to-seedling transmission of *Xanthomonas citri* pv. *malvacearum*, the causal agent of cotton bacterial blight

**DOI:** 10.1101/2021.02.09.430464

**Authors:** Jovana Mijatović, Paul M. Severns, Robert C. Kemerait, Ron R. Walcott, Brian H. Kvitko

## Abstract

Cotton bacterial blight (CBB) was a major disease of cotton in the United States in the early part of the 20^th^ century. The recent reemergence of CBB, caused by *Xanthomonas citri* pv. *malvacearum* (*Xcm*) revealed many gaps in our understanding of this important disease. In this study, we employed a field isolate of *Xcm* from Georgia USA (WT) to generate a non-pathogenic, *hrcV* mutant lacking a functional Type III Secretion System (T3SS-). We tagged the WT and T3SS- strains with an auto-bioluminescent Tn*7* reporter and compared colonization patterns of susceptible and resistant cotton seedlings using macroscopic image analysis and bacterial load enumeration. Wildtype and T3SS- *Xcm* strains colonized cotton cotyledons of resistant and susceptible cotton cultivars. However, *Xcm* populations were significantly higher in susceptible seedlings inoculated with the WT strain. Additionally, WT and T3SS- *Xcm* strains systemically colonized true leaves, although at different rates. Finally, we observed that seed-to-seedling transmission of *Xcm* may involve systemic spread through the vascular tissue of cotton plants. These findings yield novel insights into potential *Xcm* reservoirs for CBB outbreaks.

## INTRODUCTION

Cotton bacterial blight (CBB) is one of the most devastating bacterial diseases of cotton (*Gossypium* spp.)(Delannoy E. 2005). The disease is caused by the Gram-negative bacterium *Xanthomonas citri* pv. *malvacearum* (*Xcm*), and in the early and mid-20^th^ century it caused severe cotton yield losses (Hillocks 1992). For over 50 years, employed management strategies kept CBB at bay, however from 2011 *Xcm* reemerged in cotton fields (Phillips A.Z. 2017; Wang X. 2018; Rothrock C.S. 2012). Typical symptoms seen in the fields include seedling blight, angular leaf spot, vein blight, black arm, boll rot and premature boll shed (Rothrock C.S. 2015).

*Xanthomonas citri* pv. *malvacearum* (*Xcm*) is a seedborne pathogen that can survive on cotton lint on the seed surface and be transmitted to emerging seedlings (Verma 1986). However, there is some controversy about whether the pathogen is truly seed-transmitted and the significance of seedborne bacteria as a source of primary inoculum for CBB epidemics (Wang X. 2018; Innes 1983; Verma 1986; Rolfs 1915). For a pathogen to be seed transmitted, it must have the ability to gain access to the seed, survive harvest and storage, and infect and colonize emerging seedling tissues.

Within the *Xanthomonas* genus, there are many examples of seed-transmitted bacteria including *X. phaseoli* on beans (Cafati C.R. 1980), *X. manihotis* on cassava (Fritz E. 1980), *X. campestris* pv. *campestris* on cabbage (Cook A.A. 1952). Verma (1986) reported that *Xcm* could not be recovered from cotton seeds (Verma 1986). However, using a recently developed quantitative real-time polymerase chain reaction (qPCR) assay, Wang *et al* (2018) detected the pathogen in cotton seed macerate (Wang X. 2018). Although the pathogen is seed-associated, there is still some conjecture regarding the precise location of the pathogen in/on cotton seeds: internal or external to the seed coat.

Besides the localization of *Xcm* in cotton seed tissues, another unknown is whether *Xcm* is restricted to the non-vascular mesophyll tissue or if it has the ability to move through the plant via its vascular elements. Research conducted in 1956 suggested that *Xcm* colonized vascular tissues based on patterns of CBB symptom development (Wickens 1956). However, no work was subsequently done to confirm vascular tissue invasion by *Xcm*. Pathogens in the *Xanthomonas* genus display a high degree of variability in their tissue-specific invasion (Gluck-Thaler E. 2020). Plant leaf mesophyll tissue is often the target of plant pathogens; however, many destructive plant pathogens colonize xylem vessels. For example, in the case of rice there are two pathovars of *X. oryzae* that differ in their tissue localization, with *X. oryzae* pv. *oryzicola* being limited to the mesophyll and *X. oryzae* pv. *oryzae* colonizing rice xylem elements (Nino-Liu D.O. 2006). Other xylem-inhabiting Xanthomonads include the causal agents of black rot of crucifers (*X. campestris* pv. *campestris*)(Bretschneider K.E. 2004) and sugarcane leaf scald disease (*X. albilineans*) (Mensi I. 2014).

To confer disease, most Gram-negative plant pathogenic bacteria utilize a Type III Secretion System (T3SS) (Cornelis G.R.. 2000). This pathogenicity factor is used to deliver a cohort of effector proteins to interfere with the host’s immune responses or induce changes to create a conducive environment for the bacterium’s proliferation (Gala’n JE. 1999; Ghosh 2004). In the plant host, effectors or their modified targets, may be recognized by resistance proteins that activate immune reactions (Jones J.D. 2006). The ability of *Xcm* to invade different cotton tissues and the importance of its T3SS in this process can now be investigated using contemporary molecular techniques.

The majority of the research on CBB epidemiology and management was conducted prior to the deployment of resistant cotton cultivars in the early 1970s (Zhang J. 2020). For over five decades CBB was managed by acid delinting seed and by the development and deployment of resistant cotton cultivars (Zhang J. 2020; Wang X. 2018). Acid delinting chemically removes lint from cotton seed surfaces and has been used for commercial seed sanitation. However, recent research showed that acid delinting does not eliminate *Xcm* from infested cotton seeds (Brinkerhoff L.A. 1964; Wang X. 2018) but only removes bacteria from the seeds’ surface. The recent reemergence of CBB created a puzzling scenario for cotton growers and researchers.

Phillips *et al*. (2017) proffered possible explanations like the introduction of a new virulent strain, a breakdown in the cotton host resistance and the potential shift towards disease conducive environmental conditions (Phillips A.Z. 2017). They concluded that an increasing acreage planted with a susceptible cotton variety was the most likely explanation for the CBB reemergence (Phillips A.Z. 2017). However, their conclusions do not explain how the disease reemerged so suddenly, suggesting gaps in the current body of literature regarding this disease.

Here, we used molecular approaches to determine the patterns of *Xcm* colonization in seedling tissues of resistant and susceptible cotton cultivars. We also used the T3SS mutant to investigate the roles of the T3SS and effector delivery in the initial stages of seedling colonization by *Xcm*. In addition, we investigated the localization of *Xcm* in cotton seedlings after seed infection.

## MATERIALS AND METHODS

### Isolation and identification of *Xanthomonas citri* pv. *malvacearum*

*Xanthomonas citri* pv. *malvacearum* strains were isolated from CBB infected cotton leaves and bolls from outbreaks that occurred in 2017. Tissue samples were macerated in sterilized H_2_O and 100 µL aliquots of macerate were plated on Nutrient Broth Yeast agar media (8 g Nutrient broth, 2 g yeast extract, 2 g K_2_HPO_4_, 0.5 g KH_2_PO_4_, 5 g 10% (w/v) glucose solution, 0.25 g MgSO_4_, 15 g Agar and 940 ml distilled water) for 48 h at 28□. Yellow colonies were isolated and used to generate pure cultures. Overnight Lysogeny Broth (LB) (10 g tryptone, 5 g yeast extract, 10 g NaCl, pH 7.5) cultures (2 mL) were made from single colonies of putative *Xcm* strains and cell suspensions (optical density (OD) = 0.3) were generated in MilliQ ultrapure H_2_O using a spectrophotometer (Eppendorf BioSpectrometer basic, Hamburg, Germany). All strains were syringe-inoculated into 2-week-old susceptible cotton cultivar, Deltapine 1747NR B2XF (DP1747NR B2XF) (Bayer, Scott, MS) cotyledons. An *Xcm* field strain, *Xcm* 92-302, from a CBB outbreak in Georgia in 1992 was used as a positive control and *X. campestris* pv. *campestris* (strain 85-10) was used as a negative control. Strains that induced water-soaked lesions on cotton cotyledons two weeks after inoculation were confirmed to be members of the *Xanthomonas* genus by amplifying and sequencing the 16S rRNA (Eurofins, Louisville, KY, USA) as described (Barry T. 1991; Schaad N.W. 2005; Showmaker K.C. 2017). A hypersensitive response (HR) assay on tobacco leaves was conducted as previously described (Schaad N.W. 2001). However, cell death was not consistently observed by 48 h after inoculation. The identity of the *Xcm* strains was finally confirmed by PCR assay with primers described by Wang et al. (Wang X. 2018). Spontaneous rifampicin (Rif)-resistant strains of these field isolates were made by growing wildtype strains on LB agar plates amended with Rif (30 µg/ml) for 48 h. Rif-resistant *Xcm* strains were stored long-term at -80□ in 15% glycerol solution.

### Generation of a T3SS- strain of *Xcm*

To make a non-pathogenic, T3SS- strain, the *hrcV* gene was targeted based on the genome sequence of *Xcm* strain MSCT1 (National Center for Biotechnology Information (NCBI) accession number GCF_001719155.1) and a strategy was devised based on the modified allelic exchange approach (Kvitko B.H. 2011). Flanking sequences of the *hrcV* gene from the *Xcm* genome (NBCI GenBank: CP023159.1) were selected, and their merging would lead to deletion of a 771bp fragment. Gateway BP cloning *att*B1 and *att*B2 sites were added to the 5’ ends of both flanks, and the construct was synthesized by GeneUniversal (Newark, DE). The deletion construct was generated by Gateway BP-based cloning into pDONR1K18MS (Addgene plasmid #726444) and confirmed by digesting the plasmid with the restriction endonuclease *BsrG*I. The plasmid was sequenced using M13 primers (Eurofins, Louisville, KY). The Δ*hrcV* carrying plasmid was inserted into *E. coli* RHO5 by electroporation, and then mated into *Xcm* 4.02 WT by conjugation at a 1:5 (donor: recipient) ratio. The *hrcV* gene deletion in *Xcm* was confirmed by a pathogenicity test on cotton cotyledons as follows. Overnight LB broth cultures were made from single colonies of putative *Xcm* 4.02 Δ*hrcV* and *Xcm* 4.02 WT. Cell suspensions were adjusted to a concentration of ∼10^8^ CFU/ml (OD_600_ of 0.3) in MilliQ ultrapure H_2_O. Cell suspensions of all strains were syringe-inoculated into cotyledons of 2-week-old susceptible cotton (cultivar DP 1747NR B2XF) seedlings. Seedlings were observed for water-soaking symptoms for 2 weeks. Strains that did not induce water-soaked lesions were confirmed by PCR (Forward:5’- CATCACACCACCACCCTG-3’, Reverse:5’-GTCGATGAACTCGGTCGC-3’) and sequencing (Forward:5’-GTTGCAGGAAGCTGGAG-3’, Reverse:5’-CAGGTTGAATTGGGCGAC-3’) as a Δ*hrcV* deletion mutant. The *Xcm* 4.02 Δ*hrcV* strain was tested again for pathogenicity, as described above, to confirm the phenotype.

### Generation of auto-bioluminescent *Xcm* strains

Both *Xcm* 4.02 WT and *Xcm* 4.02 Δ*hrcV* were tagged with the auto-bioluminescent operon from *Photorhabdus luminescens* (*luxCDABE*) on a Tn*7* transposon as described previously (Bruckbauer S.T. 2015; Stice S.P. 2020). Donor, helper, and recipient strains were mixed in LB broth at 1:1:5 ratios, respectively, and a 10 µL aliquot of cell mixture was incubated on a sterilized nitrocellulose membrane on a LB plate supplemented with diaminopimelic acid (DAP) (400 µg/ml) at 28□. The mating mix was then grown on a LB agar plate amended with kanamycin and colonies were selected for the auto-bioluminescence phenotype. Auto-bioluminescence was detected with a sensitive charge-coupled-device (CCD) camera (Analytic Jena UVP ChemStudio, Upland, CA).

### Plant growth conditions

Cotton seeds were sown directly in soil (Farfard 3B, Sungro, Agawam, MA) in 9 cm pots, one or two seeds per pot. Pots were incubated in a growth chamber with 12 h day at 26□ and 12 h night at 23□.

### Determining auto-bioluminescence detection limits

From a single colony of *Xcm* 4.02 WT Tn*7*LUX, a culture was generated on LB agar amended with kanamycin (50 µg/mL) and incubated for 48 h at 28□. Cells were then harvested and resuspended in 10 ml sterilized H_2_O. The concentration of the cell suspension was adjusted to ∼10^8^ CFU/mL spectrophotometrically, and seven ten-fold serial dilutions were made in H_2_O. The dilution of each concentration was plated and after incubation the numbers of colonies were counted to determine concentration of the initial cell suspension. Each ten-fold serially diluted cell suspension was syringe-inoculated into 2-week-old cotton (cultivar DP 1747NR B2XF) cotyledons (4 cotyledons per inoculum level). Cotyledons were then dried with a Kimtech wipe (Kimberly-Clark, Irving, TX) and imaged for auto-bioluminescence with a CCD camera with a 2-minute exposure and no light. To determine the bacterial concentration in cotton cotyledon tissues, 3 leaf discs (∼0.1 cm radius) were collected from each inoculated cotyledon with the lowest visible and no auto-bioluminescence. The leaf discs were collected using biopsy punches (Integra Life Sciences, Plainsboro, USA) and macerated in 100 µL MilliQ H_2_O two times for 1 min at 1,750 Hz, using a GenoGrinder (SPEX SamplePrep, Metuchen, NJ). The macerate was then ten-fold serially diluted and plated on LB agar media supplemented with 50 µg/ml kanamycin. Bacterial populations were determined after incubation for two days at 28□. This experiment was repeated 3 times with two plants (4 cotyledons) per trial.

### Seed inoculation

Fungicidal seed treatments were removed from the seeds of the susceptible (DP1747NR B2XF) and resistant (PHY 480 W3FE) cotton cultivars by spraying with 70% ethanol followed by washing in sterilized H_2_O for 2 min. Seeds were then air-dried for 5 min and inoculated by submerging in 25 mL of cell suspension for 20 min. Inoculum (∼10^8^ CFU/mL) was made by suspending cells from 48 h agar plate cultures of *Xcm* 4.02 WT Tn*7*LUX and *Xcm* 4.02 Δ*hrcV* Tn*7*LUX in sterilized H_2_O. After 20 min, the inoculum was decanted and sterilized in 10% (v/v) bleach solution. Seeds were germinated for 48 h in the dark in 6 layers of sterilized cheese cloth (∼100 cm^2^) with 15 mL of sterilized H_2_O in a sterilized glass Petri dish. Germinated seeds were then planted in moist soil (Farfard 3B, Sungro, Agawam, MA) at ∼1.5 cm depth, and grown as previously described.

### Seedling sampling

Two weeks after planting, the seedlings that developed from inoculated cotton seeds were collected and imaged with a CCD camera. For each seedling, three images were taken with the following conditions: color, grayscale (70% aperture), and no light with 2 min of exposure time (100% aperture). Grayscale and 2-min exposure images were overlayed in Adobe Photoshop CC 2020. All images had the same contrast. Details regarding image processing are described in the supplemental material. Each seedling was separated into three parts: cotyledons and their petioles, stem, and all tissue starting from the first true leaf with its petiole. Each seedling component was separately weighed and macerated in 1-2 ml of sterilized H_2_O (determined based on the weight of the sample) with mortar and pestle. Tissue macerate was then ten-fold serially diluted to determine bioluminescent *Xcm* CFU per gram of cotton tissue. This experiment was repeated 3 times and for each experiment, 50 seeds were inoculated and planted.

### Statistical analysis

R-studio (RStudio 2020) was used for statistical analysis. Because the absence of colony growth generated a relatively large percentage of zeros (very often greater than 50% depending on the plant part, cultivar and treatment) in the data set and their occurrence was frequent enough to cause the underlying data to be significantly skewed, we (Log10+1) transformed the bacterial count data to improve normality, correctly apply a log transformation (e.g., log 0 = undefined), and preserve the biological signal associated with bacteria relative abundance. After transformation, the data remained non-normally distributed and unsuitable for a parametric statistical test of treatment differences based on means, thus a Kruskal-Wallis analysis of variance (ANOVA) was conducted to determine whether the median differences in the bacterial population size was statistically different between treatments and tissue types. For the same reasons, we applied a Wilcoxon Signed-Rank test to determine whether there was a significant difference in the recovered bacterial loads between treatments within tissue type (cotyledons, stems or true leaves). We also performed a *Z*-test to determine if the percentages of recovery and auto-bioluminescence differed between treatments (Table 1.) and replicated experiments (n = 3 experiments) for each tissue type.

**TABLE 1.** Analysis of percent recovery differences between experimental replicates of seed inoculated cotton seedlings.

| Cotton cultivar | Treatment | Plant part | Replicate | Number of plants |  | z score |  |  |
| --- | --- | --- | --- | --- | --- | --- | --- | --- |
|  |  |  |  | Analyzed | Recovered | I vs II | II vs III | I vs III |
| Susceptible<sup>x</sup> | *Xcm* 4.02<br>WT<br>Tn7LUX | Cotyledons | I | 22 | 21 | z= -1.1193<br>p= 0.26272 | z= -1.3579<br>p=0.17384 | z= 0.2802<br>p=0.77948 |
|  |  |  | II | 27 | 27 |  |  |  |
|  |  |  | III | 15 | 14 |  |  |  |
|  |  | Stems | I | 16 | 4 | z= 0.509<br>p= 0.61006 | z= 1.0059<br>p= 0.3125 | z= 1.3869<br>p=0.16452 |
|  |  |  | II | 22 | 4 |  |  |  |
|  |  |  | III | 15 | 1 |  |  |  |
|  |  | True leaves | I | 22 | 9 | z= -3.2386<sup>y</sup><br>p= 0.0012 | z= -3.7924<br>p= 0.00016 | z= -0.891<br>p=0.37346 |
|  |  |  | II | 27 | 23 |  |  |  |
|  |  |  | III | 15 | 4 |  |  |  |
|  | *Xcm* 4.02<br>*ΔhrcV*<br>Tn7LUX | Cotyledons | I | 15 | 14 | z= -1.2549<br>p= 0.2113 | z= -1.2549<br>p= 0.2113 | z= 0<br>p= 1 |
|  |  |  | II | 23 | 23 |  |  |  |
|  |  |  | III | 15 | 14 |  |  |  |
|  |  | Stems | I | 8 | 1 | z= 1.5297<br>p= 0.12602 | z= 1.1124<br>p= 0.267 | z= -0.4729<br>p= 0.63836 |
|  |  |  | II | 18 | 0 |  |  |  |
|  |  |  | III | 15 | 1 |  |  |  |
|  |  | True leaves | I | 15 | 4 | z= 0.1952<br>p= 0.26272 | z= 0.1952<br>p= 0.26272 | z= 0<br>p= 1 |
|  |  |  | II | 21 | 5 |  |  |  |
|  |  |  | III | 15 | 4 |  |  |  |
| Resistant | *Xcm* 4.02<br>WT<br>Tn7LUX | Cotyledons | I | 18 | 15 | z= -0.607<br>p= 0.54186 | z= -1.4152<br>p= 0.1556 | z= -1.8564<br>p= 0.6288 |
|  |  |  | II | 20 | 18 |  |  |  |
|  |  |  | III | 19 | 19 |  |  |  |
|  |  | Stems | I | 8 | 0 | z= -1.3416<br>p= 0.18024 | z= 0<br>p= 1 | z= -1.3565<br>p= 0.17384 |
|  |  |  | II | 10 | 2 |  |  |  |
|  |  |  | III | 15 | 3 |  |  |  |
|  |  | True leaves | I | 15 | 4 | z= -0.7416<br>p= 0.4593 | z= 1.1864<br>p= 0.23404 | z= 0.3832<br>p= 0.70394 |
|  |  |  | II | 18 | 7 |  |  |  |
|  |  |  | III | 19 | 4 |  |  |  |
|  | *Xcm* 4.02<br>*ΔhrcV*<br>Tn7LUX | Cotyledons | I | 16 | 16 | NA | NA | NA |
|  |  |  | II | 15 | 15 |  |  |  |
|  |  |  | III | 18 | 18 |  |  |  |
|  |  | Stems | I | 8 | 0 | NA | NA | NA |
|  |  |  | II | 15 | 0 |  |  |  |
|  |  |  | III | 15 | 0 |  |  |  |
|  |  | True leaves | I | 15 | 14 | z= -1.8194<br>p= 0.06876 | z= -1.0029<br>p= 0.31732 | z= 1.0499<br>p= 0.29372 |
|  |  |  | II | 16 | 0 |  |  |  |
|  |  |  | III | 16 | 3 |  |  |  |

<sup>x</sup> Susceptible cotton variety is Delta Pine 1747 and the resistant is Phytogen 480 W3FE
<sup>y</sup> the difference in the proportions is statistically significant (p<0.01)

### Determining localization of *Xcm* in cotton tissue

Cell suspensions were made from a 48 h plate culture of *Xcm* 4.02 WT Tn*7*LUX and cotton seeds were inoculated as described above. Twelve seeds (2 seeds each in 6 pots) were planted in soil for each inoculum level. After full expansion of the first true leaf, seedlings were collected. At this time, non-seed inoculated controls were sprayed with *Xcm* 4.02 WT Tn*7*LUX (OD_600_ = 0.3). After air-drying, surface auto-bioluminescence was confirmed by imaging with a CCD camera with a 2-min exposure time and no light. All seedlings were then surface sterilized by submerging in 70% ethanol for 1 min, with 3 subsequent washes in sterilized H_2_O. Seedlings were then dried and imaged again to determine presence and pattern of auto-bioluminescence with a CCD camera. This experiment was repeated three times.

### Scanning electron microscopy (SEM) of cotton petioles

Cotton cotyledon petioles originating from seeds inoculated with *Xcm* 4.02 WT Tn*7*LUX and non-inoculated seeds, were cut into 2 x 0.75 cm sections and placed in 2% glutaraldehyde/0.1M PBS, pH 7.2 at room temperature for 24-48 h. The samples were then rinsed 3 times for 15 min each with PBS and placed in 1% OsO4/PBS buffer for 1 h at room temperature. Samples were rinsed 3x for 15 min each with deionized H_2_O and then dehydrated with an ethanol series (25%, 50%, 75%, 85%, 95%, and three changes of 100%) for 10 min each. The samples were then transitioned into hexamethyldisilizane (HMDS) in 30 min changes of 50% ethanol/HMDS and then three changes of 100% HMDS. The samples were then air-dried in a fume hood overnight. The samples were then mounted on aluminum stubs (Electron Microscopy Sciences, Hatfield PA) with carbon adhesive and sputter coated with approximately 250 nm thick gold (Structure Probe Inc., West Chester, PA). Micrographs were obtained on an FEI/Thermo Fisher Scientific Teneo FE-SEM at 10kV. (Thermo Fisher Sci, Hillsboro, OR) at the UGA electron microscopy laboratory (Athens GA).

## RESULTS

### Isolation of *Xcm* 4.02 and generation of the T3SS- mutant

*Xcm* 4.02 was recovered from a cotton boll (cv. DP 1747NR B2XF) displaying CBB symptoms from Colquitt County, Georgia, USA. Colonies on NBY media were yellow, round, and slightly mucoid, and were confirmed to be *Xcm* by induction of CBB symptoms on cotton cotyledons and by a previously described species-specific PCR assay (Wang X. 2018). A T3SS- strain of *Xcm* 4.02 was made by allelic exchange-based deletion of a *hrcV* gene region. *Xcm* 4.02 Δ*hrcV* was confirmed by sequencing of the deleted region and the inability to induce water-soaking symptoms on cotton cotyledons after syringe infiltration (**Supplemental Figure 1.**).

### Auto-bioluminescence tagging and detection limit

Using a Tn*7*LUX auto-bioluminescent reporter (Stice S.P. 2020; Bruckbauer S.T. 2015) we observed auto-bioluminescence with 10^8^ CFU/ml and 10^7^ CFU/ml initial inoculum concentrations of *Xcm* (**Figure 1**). Auto-bioluminescence was not observed at an inoculum concentration of 10^6^ CFU/ml. The bacterial concentration at which auto-bioluminescence was not detected was 1.4 x 10^4^, and 1.9 x 10^5^ at which auto-bioluminescence was visible. The detection limit was 10^5^ CFU/cm^2^ of cotton cotyledon tissue when imaged with a 2 min exposure time (**Figure 1**).

**Figure 1.**
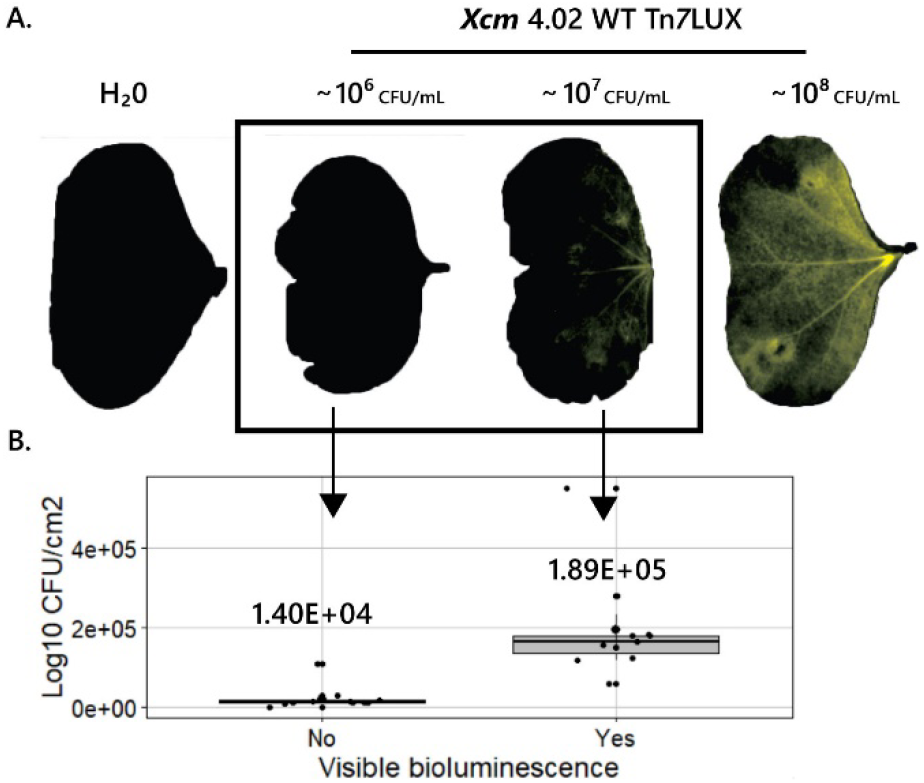
Determining visual detection limits of auto-bioluminescent bacteria *Xanthomonas citri* pv. *malvacearum* (*Xcm*) in cotton cotyledons. Auto-bioluminescence of bacteria in cotton cotyledons from 3 representative increasing inoculum doses of *Xcm* 4.02 WT Tn*7*Lux tagged bacteria compared to the water-inoculated control. Grayscale and 2-minute exposure images are overlayed with false coloring. Auto-bioluminescence is visible in yellow. The detection limit is between the highest non-visible and lowest visible Log_10_ CFU/cm^2^ (colony forming units) of cotton cotyledon tissue. The values represented are the means of 13 and 11 replicates within one experiment. Each experiment was repeated 3 times with similar results.

### Determining patterns of *Xcm* colonization after seed-to-seedling transmission

Two weeks after transplanting, cotton seedlings were analyzed for symptom development, bacterial auto-bioluminescence, and bacterial load. In 14.34% - 35.06% of cotyledons (depending on the treatment and cotton cultivar) symptoms consistent with CBB were observed, regardless of the cotton genetic background (**Supplemental Figure 2.**, Supplemental **Table 1**.). In addition, 14.34% of the water control resistant cotton plants developed CBB symptoms, although no bacteria were recovered from those plants. In 30.19% - 77.01% seedlings (depending on the treatment and cotton cultivar) we observed damage on cotyledons that is associated with partial seed coat (testa) detachment. The difference between CBB-like symptoms and seed coat damage was not clear in some cases. No CBB-like symptoms were present on stems or true leaves at the time of tissue sampling.

Cotton seedling colonization by *Xcm* 4.02 WT Tn*7*LUX and *Xcm* 4.02 Δ*hrcV* Tn*7*LUX was analyzed two weeks after seed germination. Bacteria were only detectable in susceptible cotton cotyledons developed from seeds inoculated with the *Xcm* 4.02 WT Tn*7*LUX by auto-bioluminescence visualization (**Figure 2A**). Only on one occasion (out of 64 total) were bacteria detected by auto-bioluminescence in true leaves of a susceptible cotton cultivar originating from a *Xcm* 4.02 WT Tn*7*LUX-inoculated seed (**Figure 2B**). *Xcm* populations in seedlings of the resistant cotton cultivar did not reach levels that allowed auto-bioluminescent visualization.

**Figure 2.**
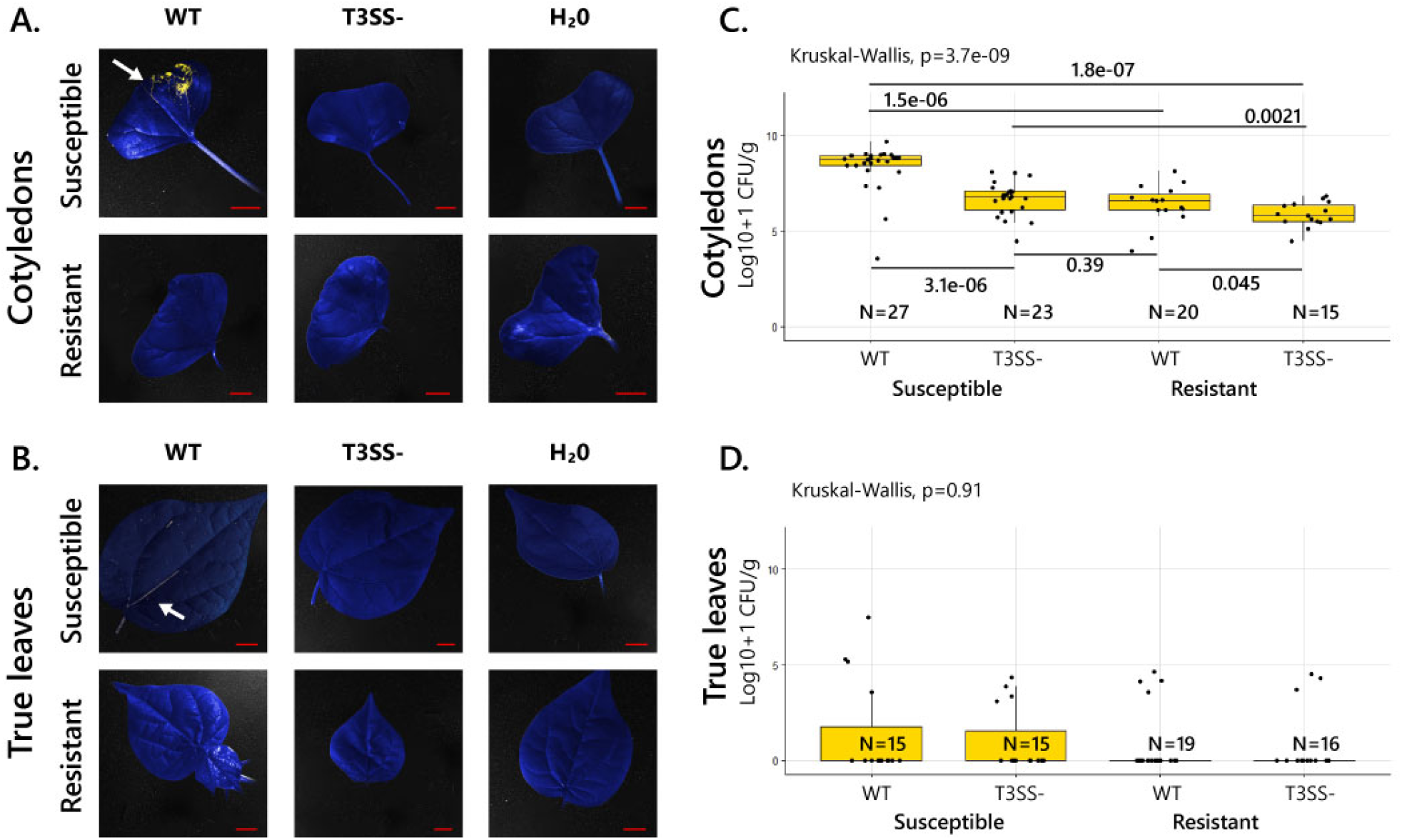
Patterns of *Xanthomonas citri* pv. *malvacearum* (*Xcm*) colonization in cotton seedling tissue after seed-to-seedling transmission. (*A)* Colonization of susceptible and resistant cotton cotyledons and (B) true leaves by *Xcm* 4.02 WT Tn*7*LUX (WT) and *Xcm* 4.02 Δ*hrcV* Tn*7*LUX (T3SS-). Grayscale image and the 2-minute exposure with no light images are overlayed. Images are representatives from 3 independent experiments. Red bars represent a 1 cm scale. Each image is cropped from a larger photo. Auto-bioluminescence is shown in yellow. (*C,D*) Comparison of the population loads of the WT and T3SS-Tn*7*Lux tagged *Xcm* strains in the susceptible and resistant cotton cultivars in Log_10_+1 CFU/g (colony forming units per gram) and incidence in cotton cotyledons (*C*) and true leaves (*D*) from one representative experimental replicate. N is the number of samples per treatment. Upper and lower quartiles are represented on all boxplots.

Cotton cotyledons were consistently colonized by *Xcm* after seed inoculation, regardless of the genetic background of the cotton cultivar or *Xcm* strain (**Figure 2C**). However, bacterial loads between all treatments and cotton cultivars were significantly different (p ≤ 0.05) (**Figure 2C**, **Supplemental figure 3B**). As expected, the *Xcm* 4.02 Δ*hrcV* Tn*7*LUX reached a lower population density compared to *Xcm* 4.02 WT Tn*7*LUX. The highest bacterial concentrations (median value LOG10+1 8.7 CFU/g) were observed in cotyledons that originated from the susceptible cotton cultivar inoculated with *Xcm* 4.02 WT Tn*7*LUX. True leaves were colonized after seedling transmission of CBB, but with significantly lower bacterial loads than those observed in cotyledons (median value LOG10+1 5.4 CFU/g) (**Figure 2D**). *Xcm* 4.02 WT Tn*7*LUX and *Xcm* 4.02 Δ*hrcV* Tn*7*LUX colonized true leaves of resistant and susceptible cotton cultivars after seedling transmission with no significant differences in population loads, but with differences in recovery rates when compared to cotton cotyledons (**Table 1**.). No auto-bioluminescence was visualized in seedling stems above the cotyledons (**Supplemental figure 3A**). There was no statistically significant difference between the *Xcm* populations in cotton stems between all treatments and cotton cultivars (**Supplemental figure 3B**).

**Figure 3.**
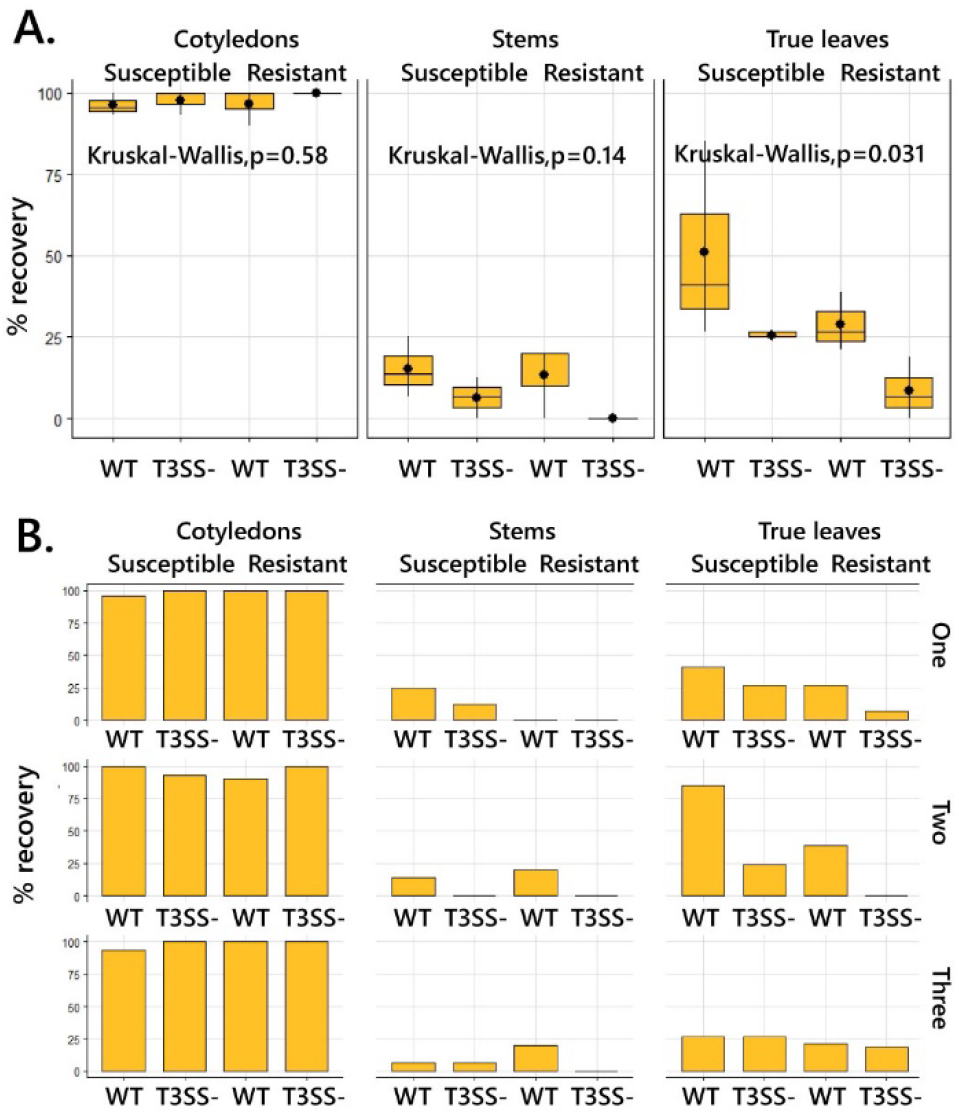
Patterns of *Xanthomonas citri* pv. *malvacearum* (*Xcm*) recovery from different seedling tissue types. *(A)* Comparison of *Xanthomonas citri* pv. *malvacearum* (*Xcm*) recovery incidences between three sampled tissue types and two cotton cultivars pulled from 3 independent experiments, inoculated with *Xcm* 4.02 WT Tn*7*LUX (WT) and *Xcm* 4.02 Δ*hrcV* Tn*7*LUX (T3SS-). (*B)* Comparison of percent recovery between experimental replicates.

Despite the lack of auto-bioluminescence detection in cotton tissues, both *Xcm* 4.02 WT Tn*7*LUX and *Xcm* 4.02 Δ*hrcV* Tn*7*LUX colonized germinated cotton seedlings after seed inoculation. Auto-bioluminescent bacteria were almost always recovered from cotton cotyledons, with overall recovery ranging from 91-100% (**Figure 3A**). Recovery of auto-bioluminescent bacteria from stems above the cotyledons ranged from 0-16.7%, with the highest recovery being recorded from stems originating from susceptible cotton seedlings inoculated with *Xcm* 4.02 WT Tn*7*LUX (**Figure 3A**). *Xcm* 4.02 WT Tn*7*LUX had the highest rate of true leaf colonization in a susceptible cotton cultivar, however the means of recovery rates in this category differed significantly (*p<0*.*01*) between experimental replicates (**Figure 3B**, **Table 1**).

### Investigating vascular presence

There was no change in the pattern of bacterial auto-bioluminescence in seed-inoculated seedlings after surface sterilization, compared to seedlings sprayed with a high concentration of *Xcm* 4.02 WT Tn*7*LUX (**Figure 4**.). The auto-bioluminescence pattern in the seedlings developed from inoculated seeds suggests systemic spread of *Xcm*. One petiole from a seed-inoculated seedling that showed auto-bioluminescence and a petiole from a seedling originating from a non-inoculated seed were subjected to scanning electron microscopy. Imaging of the xylem vessels revealed bacterial colonization in all samples of the petiole originating from the *Xcm* 4.02 WT Tn*7*LUX-inoculated seedling. Bacterial cells were not observed in the xylem of the samples originating from the petiole of the water-inoculated control (**Figure 5**.).

**Figure 4.**
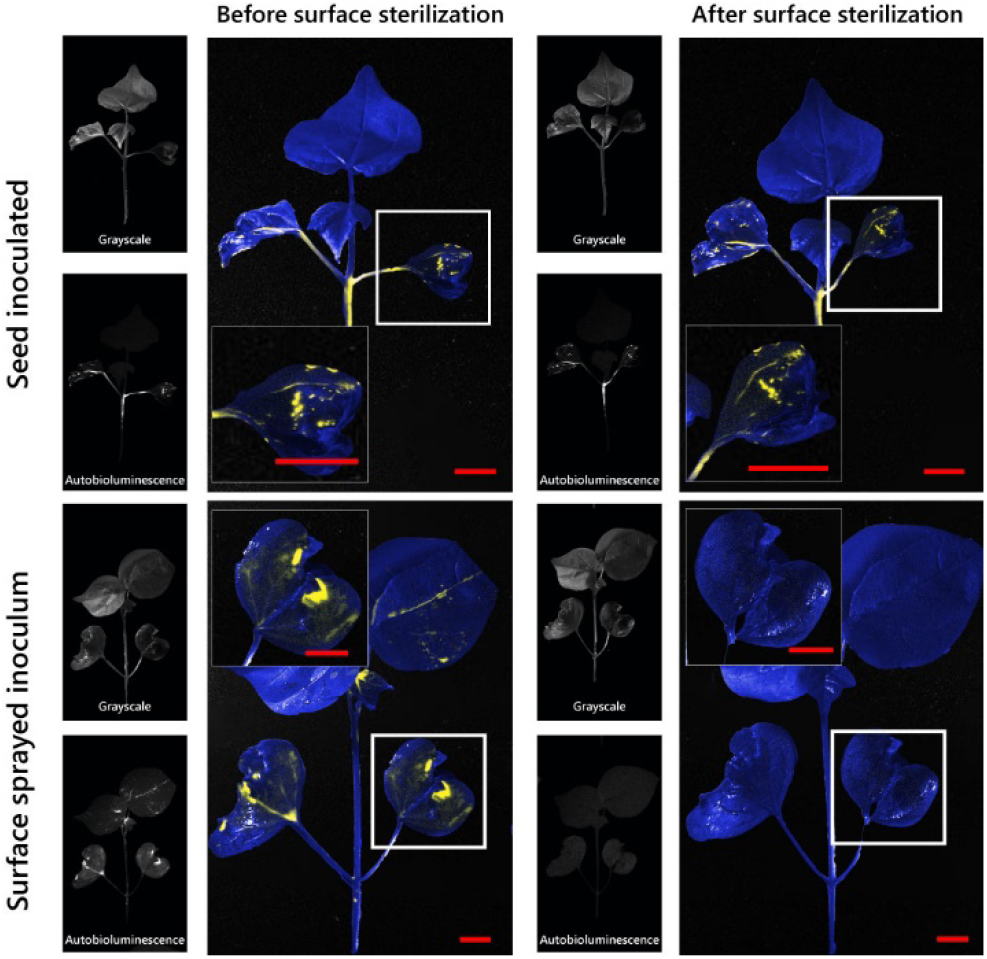
Evidence of internal cotton seedling tissue colonization by *Xanthomonas citri pv. malvacearum (Xcm)* after seed infection. Different treatments of cotton seedlings before and after surface sterilization with 70% ethanol. First and third column are the grayscale and long exposure photos of seedlings. Second and fourth columns are overlays of the grayscale and ling exposure photos of cotton seedlings that were either seed inoculated with OD_600_ of 0.3 (∼10^8^ CFU/mL) of *Xcm* 4.02 WT Tn*7*LUX or surface sprayed with high inoculum concentration of *Xcm* 4.02 WT Tn*7*LUX. Images are cropped from a larger file and are representatives of one independent experiment. Auto-bioluminescence is visible in yellow. Red bar is 1 cm.

**Figure 5.**
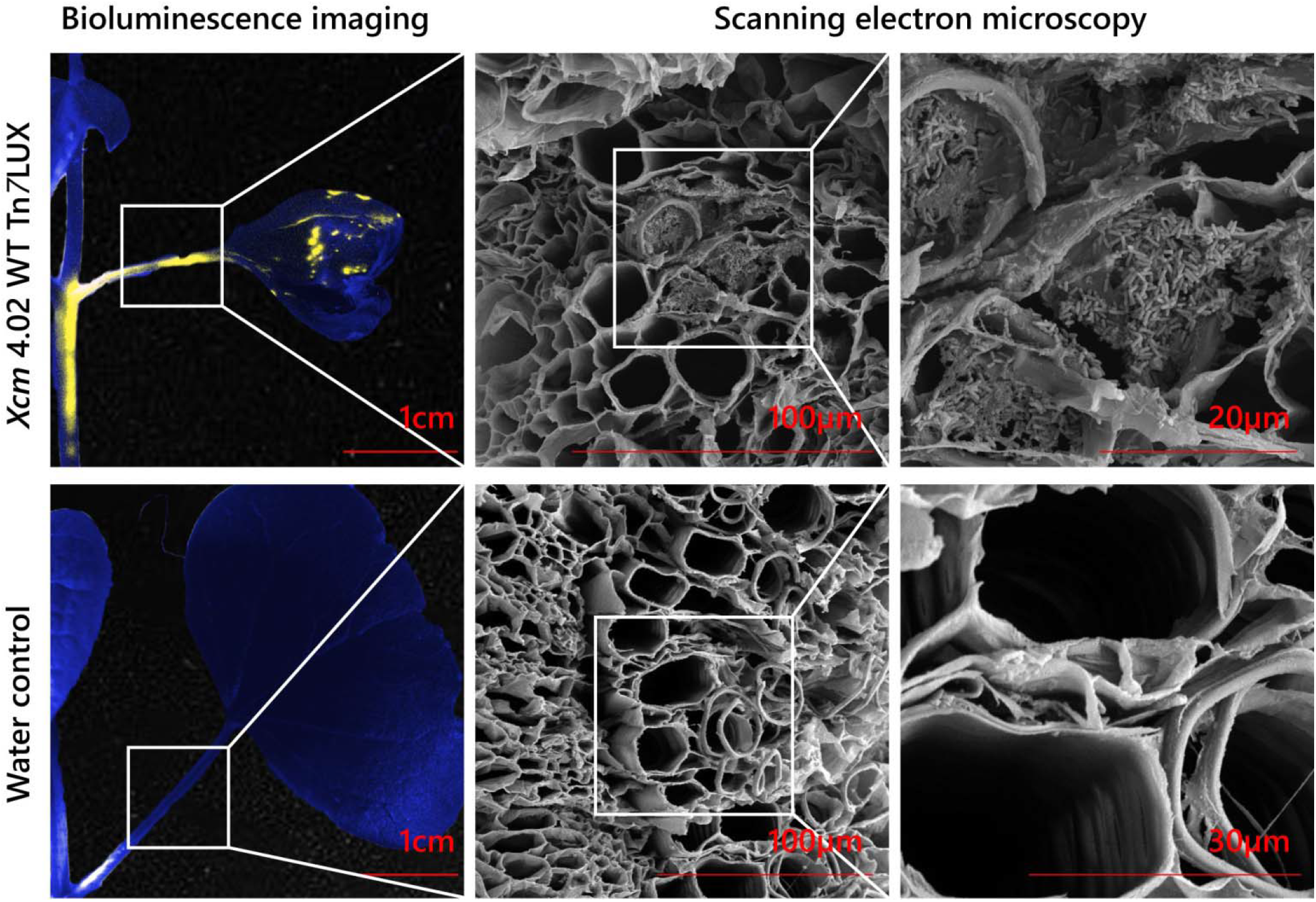
Evidence of vascular colonization of cotton seedlings by *Xanthomonas citri* pv. *malvacearum (Xcm)* after seed inoculation. Petioles of seedlings from *Xcm* 4.02 WT Tn*7*Lux and a water-inoculated seeds, 2 weeks post sprouting and after surface sterilization. First panels are image overlays to show auto-bioluminescence presence. Second two panels show results of scanning electron microscopy of the xylem vessels of the petioles. Auto-bioluminescence is visible in yellow. Red lines represent length for scale.

## DISCUSSION

Cotton, one of the most important fiber crops in the world, is under constant attack by many plant pests and pathogens (Ritchie G.L. 2007; Cox K.L. 2019; Kirkpatrick T.L. 2001). Cotton bacterial blight was first reported in 1891 by Atkinson (Atkinson 1891), and since then, research efforts have sought to limit outbreaks and yield losses. The recent CBB epidemics have instigated work to understand the genetic basis of disease reemergence (Phillips A.Z. 2017; Cox K.L. 2017; Cox K.L. 2019; Showmaker K.C. 2017; Cunnac S. 2013), improve detection methods (Wang X. 2018; Ajene I.J. 2016) and understanding characteristics of the CBB-resistant cotton cultivars (Zhang J. 2020). Phillips *et al*. (2017) posited that the underlying cause of reemergence was the increase in cultivation of CBB susceptible cotton cultivars (Phillips A.Z. 2017). Their research also revealed the lack of information regarding the CBB epidemiology that would explain how this pathogen survived over 50 years without a major CBB outbreak. We investigated aspects of CBB seed transmission and seedling colonization patterns and revealed a potential reservoir of CBB.

Currently, CBB management relies on acid delinting of seeds and use of resistant cotton cultivars (Wang X. 2018; Phillips A.Z. 2017). Although acid delinting is purported to remove *Xcm* from the seed surface, bacterial cells can survive the process (Wang X. 2018; Alexander A. S. 2012; Hunter R.E 1964). Here, we investigated the ability of *Xcm* to establish disease on cotton seedlings after seed inoculation/seedling transmission in a susceptible and a resistant cotton cultivar. Results from the current study indicate that initially, *Xcm* colonizes cotton cotyledons regardless of the genetic background of cotton or bacteria (**Figure 3A**). We also observed a significant difference in bacterial recovery from true leaves in the susceptible cotton cultivar (**Figure 3B**, **Table 1**). Since all seedlings were treated in the same way and grown in controlled conditions, the factor that contributed to higher true leaf tissue colonization in our second experimental replicate is unclear.

Wang et al. reported that discerning CBB symptoms and foliar damage caused by other factors (e.g. seed coat release) may be difficult (Wang X. 2018). In line with these findings, we found that some cotyledons developed CBB symptoms, but in many cases, the symptoms were not distinguishable from damage associated with poor seed coat detachment during seedling development (**Supplemental figure 2**, **Supplemental table 1**).

Of all of the cotton seedling tissues tested, only cotyledons were colonized with sufficient bacteria loads for auto-bioluminescence visualization (**Figure 2A**). In one case, the bacterial load was high enough to visually detect auto-bioluminescence in the true leaf of a susceptible cotton cultivar inoculated with *Xcm* 4.02 WT Tn*7*LUX (**Figure 2B**). While we documented low auto-bioluminescence visualization rates, our experiments also showed that the absence of auto-bioluminescence does not imply the absence of bacteria (**Figure 1**). Regardless of the lack of auto-bioluminescence, both *Xcm* 4.02 WT Tn*7*LUX and *Xcm* 4.02 Δ*hrcV* Tn*7*LUX colonized cotton seedlings after seed infection. Even though population loads recovered from cotyledons varied significantly depending on the cotton cultivar and bacterial genetic background (**Figure 2C**), we observed 91-100% recovery of *Xcm* strains from cotyledons (**Figure 3A**). Although we document that the *Xcm* population loads and percent recovery were lower in true leaves than those observed in cotyledons (**Figure 2C, 2D** and **3A**) at the time of sampling (about 2 weeks post-germination), it is still unknown how the population dynamics change at a later stage of cotton development.

The T3SS is necessary for disease induction by many Gram negative plant pathogenic bacteria (Alfano J. R. 1997). However, in some cases it is not utilized in the initial stages of disease (Johnson K.L. 2011; Hirano S.S. 1999; Darrasse A. 2010). Consistent with these results, our data showed no significant difference in the recovery of *Xcm* from cotton cotyledons of resistant or susceptible cotton cultivars originating from seeds inoculated with *Xcm* 4.02 WT Tn*7*LUX and *Xcm* 4.02 Δ*hrcV* Tn*7*LUX (**Figure 3A**). In addition, the recovered populations of 4.02 Δ*hrcV* Tn*7*LUX in the susceptible cotton cultivar were lower than those recovered from seeds inoculated with *Xcm* 4.02 WT Tn*7*LUX (**Figures 2C** and **2D**, **Supplemental figure 3B**). These results indicate that the initial stages of cotton seedling colonization by *Xcm* does not require a functional T3SS, even in the resistant cotton cultivar.

Using Tn*7*LUX-tagged *Xcm* strains we macroscopically visualized bacterial populations in cotton tissue. Imaging seedlings grown from inoculated seeds revealed a pattern of *Xcm* auto-bioluminescence along the cotyledon vasculature. In order to rule out potential auto-bioluminescence of surface populations, we recorded the patterns of colonization after seed infection, pre- and post-surface sterilization (**Figure 4**). Our results show no change in pattern in the bacterial auto-bioluminescence in seed-inoculated seedlings after surface sterilization, compared to seedlings sprayed with a high concentration of *Xcm* 4.02 WT Tn*7*LUX whose auto-bioluminescence pattern was lost. These results indicate that bioluminescent bacteria imaged on seed-inoculated seedlings were not removed by seedling surface sterilization and that the bacteria were present within the seedling tissue. To further investigate the hypothesis of *Xcm* being present within cotton seedling vascular tissues, we analyzed a petiole sample from a surface sterilized infected seedling by SEM. We observed bacteria in the vascular elements of the cotyledon petiole originating from the *Xcm* 4.02 WT Tn*7*LUX-inoculated seed (**Figure 5**.). Bacteria were not observed in the xylem vessels of the cotyledon petioles originating from the water-treated cotton seeds. These results are consistent with the hypothesis of *Xcm* entering the cotton vasculature and being systemically transmitted through the plant after seed infection.

Overall, this study demonstrates that cotton seedlings are colonized by *Xcm* and that *Xcm* can colonize the vasculature of cotton seedlings when infected seeds germinate. In addition, we observed tissue colonization with and without symptom development, congruent with previous work (Wang X. 2018). Together, findings from the current study add to the growing body of literature that plants colonized with *Xcm* below the threshold of symptom development may act as potential reservoirs of *Xcm* in cotton fields grown between outbreaks. However, the data do not rule out the role other inoculum sources like volunteer plants and cotton debris that may have instigated recent CBB outbreaks in cotton fields (Brinkerhoff 1970; Verma 1986). Nevertheless, our data demonstrates that *Xcm* can be seed transmitted and can colonize seedlings of a resistant cotton cultivar. While we document that *Xcm* can colonize the xylem under the tested conditions, the factors that affect possible systemic movement and spread are yet to be determined. In addition, the successful colonization of the cotton seedlings by *Xcm* 4.02 Δ*hrcV* Tn*7*LUX indicates a potential ontogenetic aspect of CBB resistance.

## Acknowledgements

We thank Shaun Stice and Amy Smith (Univ. of Georgia) for technical assistance. The authors also thank Li Yang (Univ. of Georgia) for the use of laboratory equipment. Additional thanks are extended to John Shields (Univ. of Georgia, Georgia Electron Microscopy) for his assistance with sample preparation and Scanning Electron Microscopy. Lastly, we thank the Georgia Commodity Commission for cotton for their financial support.

## Notes

### Competing Interest Statement

The authors have declared no competing interest.

