## Supplemental materials for "Elucidating the patterns of seed-to-seedling transmission of *Xanthomonas citri* pv. *malvacearum*, the causal agent of cotton bacterial blight"

**Supplemental material:**

**Processing of auto-bioluminescence images.**

Grayscale and auto-bioluminescence-captured images were processed in Adobe Photoshop CC2020 (Adobe Inc.). Both photos were opened, and the auto-bioluminescence photo was overlaid onto the grayscale photo. On the top toolbar, the Image>Mode>CYMK>Do not flatten/merge path was followed, and Layer 0 was unlocked. The Base 1 file was selected and layers Levels 1 (Black point= 35, Grey point= 1.60, White point= 215) and Gradient map 1 ( transparency at 20%) were selected and duplicated to the working file. Levels 2 (Black point= 28, Grey point= 0.89, White point= 58) and Color fill (#fff200 or C=0,Y=100, M=0, K=0) were selected and duplicated to the working file. This step was done to ensure uniform image processing. Layer order from top to bottom was: Color fill 1, Levels 2, Layer 1, Gradient map 1, Levels 1 and Layer 0. Layer 1 was selected, and clipping mask was released. Layer 1 was set as a Light color instead of Normal. Gradient map 1 and Levels 1 layers were already selected as a clipping mask. Image was saved as JPEG. The new JPEG image and the grayscale image were opened, and the new JPEG image was overlayed over the grayscale image. While only grayscale image was visible and selected, the plant tissue was selected with a quick selection tool. When done, select Layer>New>Layer via copy path was followed. The saturation layer ( Hue=236, Saturation=73, Lightness= +5, at 20% transparency) from the Base 2 file was copied to the new working file. This step was done to ensure uniform image processing. The order of layers from top to bottom was: Layer 1, Hue/Saturation 1, Layer 2, Layer 0. Select Layer 1 and click Layer>Make clipping mask and Lighten instead of Normal was selected. Final files were saved both as an Adobe Photoshop file and JPEG.

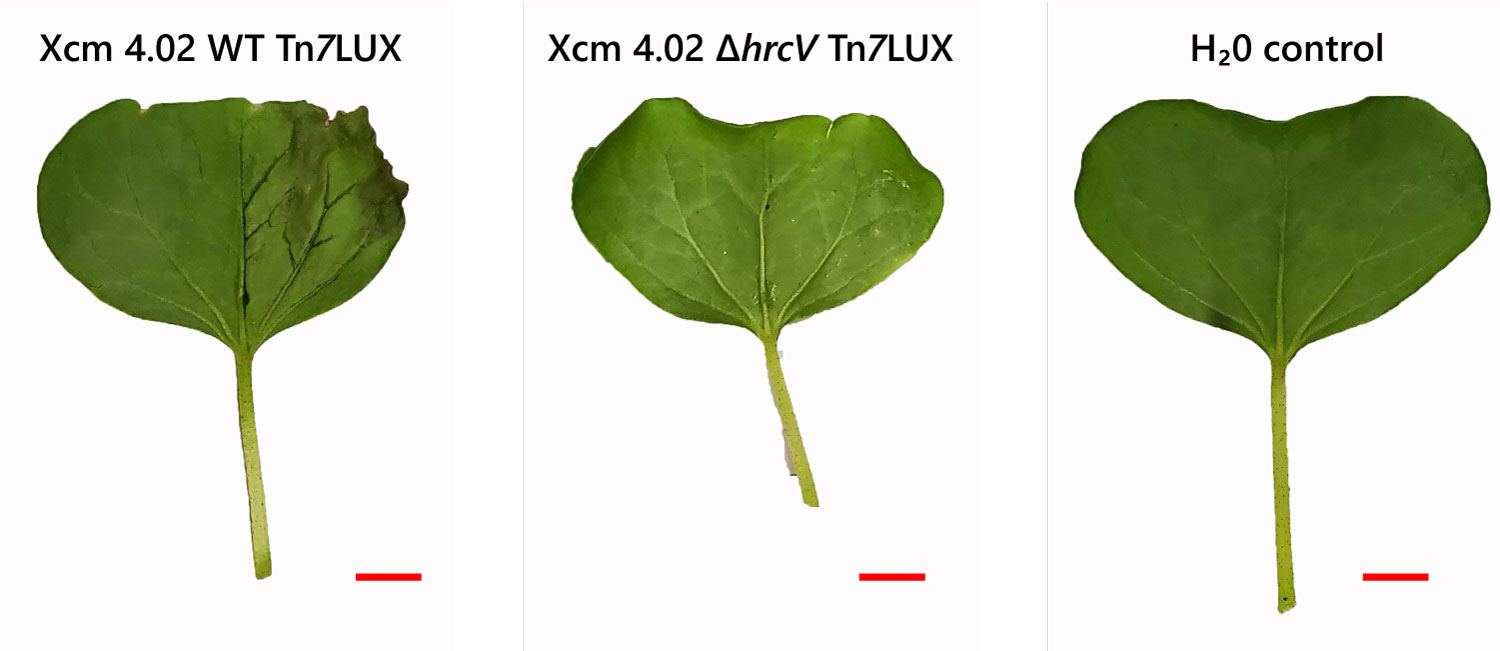

**Supplemental Figure 1. T3SS- *Xanthomonas citri* pv. *malvacearum (Xcm)*** **does not cause symptoms on cotton cotyledons.** Cotyledons of two-week-old cotton seedlings were inoculated with the *Xcm* 4.02 WT Tn*7*LUX and *Xcm* 4.02 Δ*hrcV* Tn*7*LUX strains at OD600 of 0.3 (~10^8^ CFU/ml) in sterilized H_2_O, and imaged 10 days after inoculation. Experiment was repeated 3 times with similar results and 3 representative samples are shown.

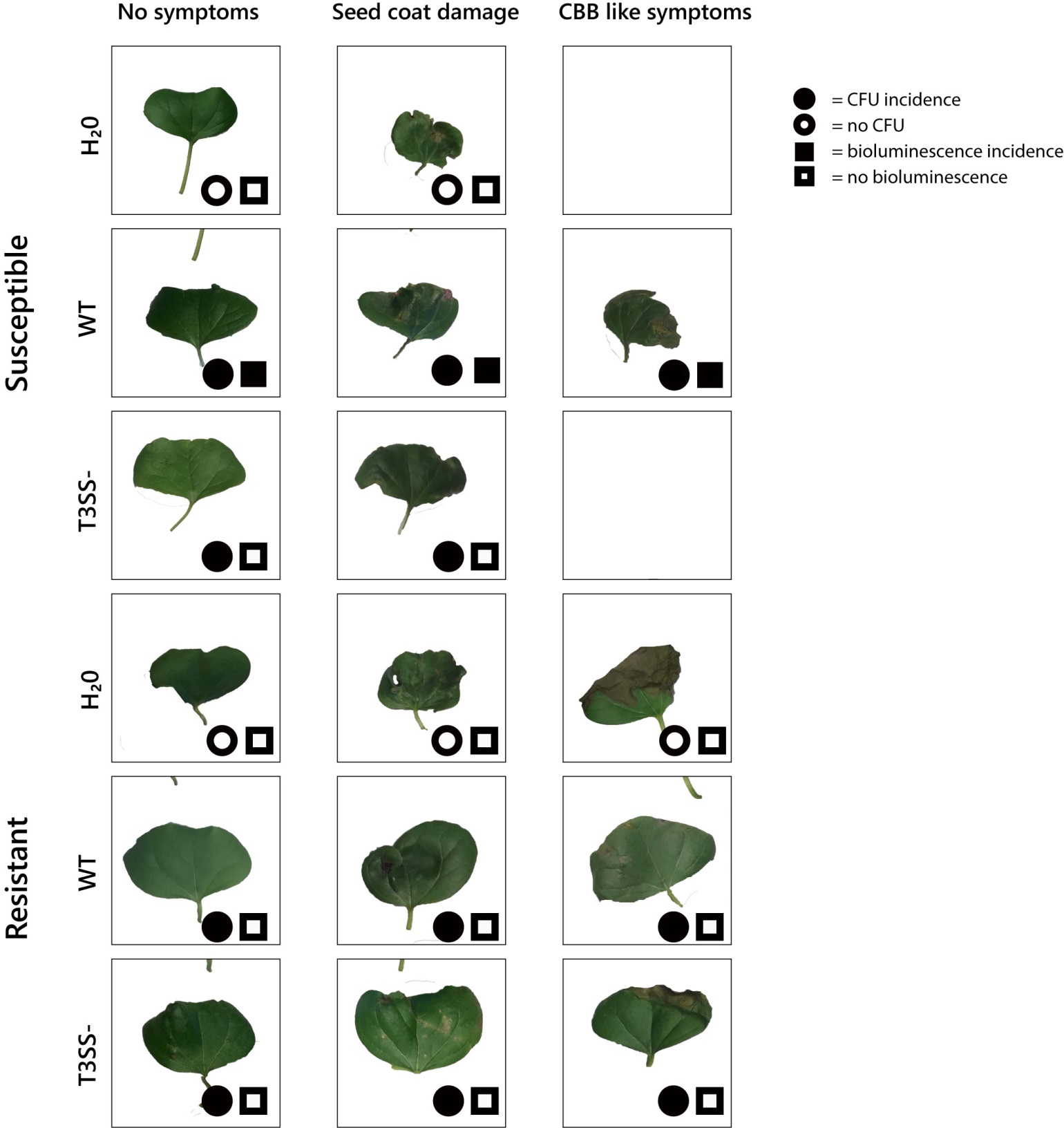

Supplemental figure 2. **Cotton cotyledons colonized by *Xanthomonas citri* pv. *malvacearum*  (*Xcm*) without cotton bacterial blight (CBB) symptom development.** *Xcm* 4.02 WT Tn*7*LUX (WT) and *Xcm* 4.02 Δ*hrcV* Tn*7*LUX (T3SS-) strains. Representative images of cotton cotyledons showing no symptoms, coat damage symptoms and CBB symptoms. Map shows the status of colony forming unit recovery and auto-bioluminescence visualization with 2-minute exposure.

Supplemental Table 1. Percent incidence of seed coat associated damage and cotton bacterial blight-like symptoms.

| Cotton line | Treatment | Replicate | seed coat associated damage % | CBB like symptom  % | total sample number |
| --- | --- | --- | --- | --- | --- |
| Susceptible cotton line | DP^x^ H_2_0 | Experiment 1 | 72.73 | 0.00 | 11 |
|  |  | Experiment 2 | 60.00 | 0.00 | 10 |
|  |  | Experiment 3 | 80.00 | 0.00 | 15 |
|  |  | Average | 70.91 | 0.00 | 36 |
|  | DP *Xcm* 4.02 WT Tn*7*LUX | Experiment 1 | 63.64 | 0.00 | 22 |
|  |  | Experiment 2 | 74.07 | 51.85 | 27 |
|  |  | Experiment 3 | 93.33 | 53.33 | 15 |
|  |  | Average | 77.01 | 35.06 | 64 |
|  | DP *Xcm* 4.02 Δ*hrcV* Tn*7*LUX | Experiment 1 | 47.83 | 0.00 | 15 |
|  |  | Experiment 2 | 73.91 | 0.00 | 23 |
|  |  | Experiment 3 | 35.29 | 0.00 | 15 |
|  |  | Average | 52.34 | 0.00 | 53 |
| Resistant cotton line | PH H_2_0 | Experiment 1 | 54.55 | 36.36 | 11 |
|  |  | Experiment 2 | 12.50 | 6.67 | 16 |
|  |  | Experiment 3 | 23.53 | 0.00 | 17 |
|  |  | Average | 30.19 | 14.34 | 44 |
|  | PH *Xcm* 4.02 WT Tn*7*LUX | Experiment 1 | 66.67 | 61.11 | 18 |
|  |  | Experiment 2 | 47.37 | 0.00 | 20 |
|  |  | Experiment 3 | 80.00 | 6.67 | 19 |
|  |  | Average | 64.68 | 22.59 | 57 |
|  | PH *Xcm* 4.02 Δ*hrcV* Tn*7*LUX | Experiment 1 | 31.25 | 25.00 | 16 |
|  |  | Experiment 2 | 80.00 | 26.67 | 15 |
|  |  | Experiment 3 | 27.78 | 11.11 | 18 |
|  |  | Average | 46.34 | 20.93 | 49 |

^x^ Susceptible cotton variety is DP1747NR B2XF (DP) and the resistant is Phytogen W3FE (PH)

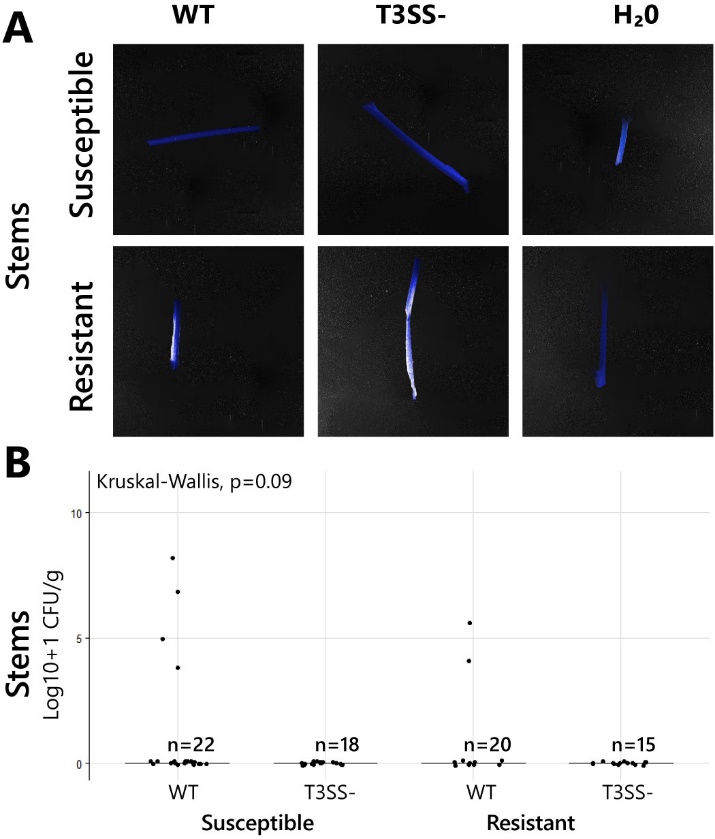

Supplemental Figure 3. **Patterns of *Xanthomonas citri* pv. *malvacearum* (*Xcm*) colonization after seed-to-seedling transmission** (A) Auto-bioluminescence is not visible on cotton seedling stems regardless off the cotton cultivar or treatment type. Auto-bioluminescence if present, would be visible in yellow. (B) Comparison of the population loads of the *Xcm* 4.02 WT Tn*7*LUX (WT) and *Xcm* 4.02 Δ*hrcV* Tn*7*LUX (T3SS-) strains in the susceptible and resistant cotton cultivars in Log10+1 CFU/g (colony forming units per gram) in cotton stems. n is the number of samples.
